# Prucalopride ameliorates delayed gastrointestinal transit and social behaviour in a mouse model of 15q duplication syndrome

**DOI:** 10.1101/2025.03.03.640221

**Authors:** Gayathri K. Balasuriya, Kota Tamada, Jun Nomura, Carla Cirillo, Toru Takumi

## Abstract

**Introduction:** Chromosome 15q duplication syndrome (Dup15q) is a neurodevelopmental disorder linked to autism spectrum disorder (ASD), involving increased copies of the 15q11.2-q13 region. About 80% of individuals with Dup15q experience gastrointestinal (GI) dysfunction, including constipation. The duplicated region encodes GABA receptor A subunits, affecting GABAergic signalling, while reduced serotonin (5-HT) levels impair neuronal activity and social behaviour in a mouse model of Dup15q (15q dup). Given the importance of GABA and serotonin in the enteric nervous system (ENS), this study investigates GI dysfunction and neurotransmission in a Dup15q mouse model.

**Methods:** Colon RNA extracts were analysed for GABA receptor subunit and serotonin-associated gene expression using quantitative PCR. Total GI transit was assessed by Carmine red dye gavage. Ex vivo colonic motility was analysed via video imaging. The GABA receptor A antagonist Bicuculline was used to assess GABAergic signalling. Prucalopride, a 5-HT4 receptor (5HT4R) agonist, was administered for six days, and its effects on GI transit and social interaction were evaluated.

**Results:** 15q dup mice exhibited elevated GABA receptor gene expression and reduced Tph2 and Htr4 expression in the colon. Total GI transit was delayed, and ex vivo colonic motility was slower and less extensive. Bicuculline further impaired colonic contractions, indicating enhanced GABAergic sensitivity. Prucalopride restored GI transit delays and improved social interaction, as evidenced by increased contact duration in social tests.

**Conclusion:** Prucalopride effectively restores GI function and improves social behaviour in 15q dup mice, demonstrating its therapeutic potential for addressing both GI dysfunction and behavioural deficits in 15q duplication syndrome.

## Introduction

Gastrointestinal (GI) dysfunction is a prevalent comorbidity in individuals with neurodevelopmental disorders, including autism spectrum disorders (ASD), significantly affecting the quality of life [1–3]. ASD, characterized by deficits in social communication, restricted interests, and repetitive behaviours, affects approximately one in 36 children aged 8 years in the USA in 2020 [4]. Genetic factors play a major role in its pathogenesis, with various susceptibility genes and chromosomal abnormalities implicated. In addition to core features, individuals with ASD often experience GI issues, such as constipation, diarrhea, and abdominal pain, affecting up to 48%-90% of individuals [1, 5]. The impact of GI dysfunction in ASD extends beyond physical discomfort; it can significantly impair quality of life, disrupt sleep patterns, interfere with educational and therapeutic interventions, and potentially exacerbate behavioural challenges such as irritability and aggression [6].

15q duplication syndrome (Dup15q), a genetic subtype of ASD resulting from duplication of the 15q11.2-q13 chromosomal region, provides a valuable model for investigating the interplay between ASD, GI dysfunction, and behaviour. Dup15q is one of the most common genetic causes of ASD, accounting for up to 0.5-3% of cases [7]. Individuals with Dup15q frequently exhibit core ASD features as well as a high prevalence of GI issues, with constipation being a particularly common complaint presented in up to 60% of the patients surveyed [8, 9]. Clinical observations in Dup15q suggest a potential link between GI dysfunction and behavioural challenges, with improvements in irritability and aggression, and report following successful treatment of GI symptoms [8]. Studying GI dysfunction in Dup15q is important, as research shows 49% of children with Dup15q experience GI symptoms before diagnosis [9].

The enteric nervous system (ENS) plays a crucial role in regulating GI motility and function. Within the ENS, neurotransmitter systems, including gamma-aminobutyric acid (GABA) and serotonin (5-hydroxytryptamine, 5-HT), are pivotal in maintaining gut homeostasis [10, 11]. The gut-brain axis, a bidirectional communication system between the GI tract and the central nervous system, is increasingly recognized as a key player in neurodevelopment and behaviour [12]. Alterations in gut motility, intestinal permeability, microbiota composition, and immune function have been implicated in the pathophysiology of ASD and other neurodevelopmental disorders [13–16]. Additionally, dysregulation of the hypothalamic-pituitary-adrenal (HPA) axis, often reflected in altered corticosterone levels in rodent models, has been linked to both GI dysfunction and behavioural abnormalities in ASD-related conditions [17]. Given that stress and neuroendocrine factors influence gut function and behaviour, we assessed corticosterone concentration as a potential biomarker of stress-related physiological alterations in 15q dup mice.

Prucalopride, a high-affinity serotonin receptor (5-HT4R) agonist approved for the treatment of chronic constipation [18, 19], offers a potential means to target both GI and behavioural symptoms in Dup15q. By enhancing GI motility and promoting propulsive contractions, Prucalopride may alleviate constipation and potentially improve related behavioural difficulties.

In this study, we utilized a well-established Dup15q mouse model (15q dup), which involves the duplication of mouse chromosome 7 corresponding to the human chromosome 15q11-13 region [20]. These 15q dup mice exhibit behavioural traits reminiscent of those observed in individuals with ASD, including deficits in social communication and behavioural inflexibility, along with reduced brain 5-HT levels, impaired synaptic plasticity, disruptions in excitatory/inhibitory balance in the somatosensory cortex and altered gut microbiome [15, 20–23]. Here, we investigated the effects of Prucalopride on GI transit, as well as its potential to improve social behaviours. This study aims to uncover the interplay between gut function, behaviour, and pharmacological intervention in this complex syndrome.

## Methods

### Animals

The Dup15q mouse model (15q dup) was generated as previously described [20] and bred for over 10 generations on a C57BL/6J background. Wild-type (WT) and 15q dup littermates were cohoused in a controlled environment with a 12-hour light/dark cycle (lights on at 6:00 am and off at 6:00 pm). All mice were provided ad libitum access to food and water. Both male and female mice were used for most experiments, while detailed colonic motility analyses were performed exclusively on male mice. All experimental procedures were approved by the Kobe University Graduate School of Medicine Animal Ethics Committee and conducted in accordance with the ARRIVE Guidelines for the reporting of animal research.

### Quantitative RT-PCR

Mice were sacrificed using isoflurane anaesthesia followed by cervical dislocation. Microdissection of colon tissue was performed to isolate the longitudinal muscle layer plus myenteric plexus preparation (LMMP) as previously described [14]. Total RNA was extracted from the LMMP using TRIZOL reagent (Thermo Fisher Scientific), and reverse transcription was carried out with SuperScript IV (Thermo Fisher Scientific, USA). Quantitative RT-PCR was conducted as described in prior studies [20], with Gapdh used as the internal housekeeping gene. Relative gene expression was calculated using the delta-delta CT method [24]. The primer sequences are listed in Supplementary Table 1.

### Histopathological assessment

#### H&E staining

For histological analysis, 4% PFA fixed paraffin embedded colon and small intestine were sectioned (10 µm thickness), mounted on glass slides and deparaffinized. Slices were stained by using the standard hematoxylin and eosin (H&E) staining method [25]. Brightfield images (X20 objective) were obtained using an Olympus slides scanner (Olympus VS120) and functions of ImageJ (NIH, USA) were used to measure parameters such as villus height, villus width, crypt depth, and muscle layer thickness.

#### Alcian Blue staining for mucus layer thickness

Alcian Blue staining was performed on mid-colon tissues (1–2 cm segments containing a faecal pellet) following an established protocol [26, 27]. Brightfield images of the stained sections were acquired using an Olympus slide scanner (VS120). The mucus layer thickness was quantified using ImageJ software (NIH, USA), and the average thickness was calculated for each sample. See the supplementary methods section for further information.

### Immunofluorescence

#### Wholemount preparation and immunofluorescence study of the ENS

Wholemount preparations of the ENS were obtained following previously established protocols [25]. Briefly, the gut tissue was opened along the mesenteric border, pinned flat with the mucosal side facing upward on a Sylgard-coated (Dow Corning, USA) petri dish, and fixed in 4% paraformaldehyde at room temperature for 80 minutes. To isolate wholemount preparations of myenteric neurons, the mucosa and circular muscle layers were carefully removed under a dissecting microscope using fine forceps. Preparations were blocked using 10% CAS-Block (Invitrogen) and 0.1% Triton X-100 (NACALAI TESQUE, INC.) solution for 30 minutes at room temperature. Following blocking, tissues were incubated overnight at 4°C with primary antibodies, followed by incubation with corresponding secondary antibodies for 2 hours at room temperature. Samples were mounted on glass slides using VECTASHIELD Mounting Medium with DAPI (Vector Laboratories). Images (20X) were acquired using a confocal microscope (FV3000). Image analysis and cell quantification were performed using ImageJ software (NIH, USA).

#### Iba1 immunofluorescence in colon cross section

Colon tissues were fixed in 4% PFA, cryoprotected in 30% sucrose, and sectioned at 10 µm. Sections were blocked with 10% CAS-Block and 0.1% Triton X-100, incubated with Iba1 primary antibody overnight, followed by secondary antibody. After mounting with VECTASHIELD Mounting Medium with DAPI, images were acquired using an Olympus slides scanner (VS120). Iba1 fluorescence intensity was quantified using ImageJ (NIH, USA).

Antibody information is listed in Supplementary Table 2.

### GI transit

#### Total GI transit

We used the Carmine red dye oral gavage method to study the total GI transit time [14]. Briefly, eight weeks old 15q dup and WT mice were fed 0.2 mL Carmine red dye (20 mg/mL, Sigma) by oral gavage and time taken to defecate a red pellet was recorded and considered as an estimate of the total GI transit time.

#### Small Intestinal Motility

Mice were orally gavaged with 0.2 mL Carmine red dye (20 mg/mL, Sigma). After 10 minutes, mice were euthanized, and the gastrointestinal tract was removed. Small intestinal transit was assessed by measuring the distance travelled by the dye front and calculating the ratio of this distance to the total small intestinal length. [14].

#### In vivo colonic motility assessment using glass bead propulsion test

8-week-old mice were first fasted for 12 hours with access to water Eight-week-old mice were fasted for 12 hours with ad libitum access to water to minimize the influence of faecal pellets in the distal colon on experimental outcomes. Animals were anaesthetised using 5% isoflurane inhalation before inserting a 3 mm diameter glass bead 2 cm into the colon using a sterile glass rod [28]. The time taken to expel the glass bead was recorded and used as an estimate of colonic motility.

#### Video imaging of the colonic motility

Ex vivo colonic motility was assessed in male mice, as the syndrome exhibits a 2:1 male bias. Additionally, our previous studies using the same technique have demonstrated strong estrus cycle-dependent effects on colonic motility, which could introduce variability in the data. Adult male WT and 15q dup mice (aged 8–10 weeks) were culled by cervical dislocation and the entire colon was dissected out to study the ex vivo colonic motility using video imaging technique as previously reported [29, 30]. The colon was equilibrated in the organ bath for 30 min before recording 4 control videos of 15 min each. Colons were then exposed to Bicuculline (10 μM, Sigma-Aldrich) and four 15 min videos were recorded. (Details in supplementary methods).

#### Prucalopride drug intervention study

The experimental design is shown in Figure 5A. Baseline GI transit and baseline assessments (weight, faecal corticosterone, social interaction) were performed. Mice received daily oral gavage of Prucalopride (1 mg/kg, Selleckchem) dissolved in 200 µl of saline or an equal volume of saline for 6 days [31, 32]. On day 6, these assessments were repeated. On day 7, GI transit was re-assessed.

#### Reciprocal social interaction test

Reciprocal social interaction between two mice was assessed in a novel cage with fresh bedding (dimensions: 30.80 × 59.37 × 22.86 cm). The paired mice were of the same genotype and sex but from different home cages and had comparable body weights. The test arena was illuminated with dim light (10 lux) to minimize stress, and the interactions were recorded using an overhead CCD camera for 10 min [21]. No aggressive behaviour has been previously reported in these mice [21]. However, if aggression occur during experiments, the effected mice will be immediately removed. The duration of social interactions, including sniffing and physical contact duration, was manually quantified.

#### Faecal corticosterone measurement

Measurements were made from the faecal samples collected from the cages during social interaction tests. Samples were immediately frozen and analysed using an ELISA kit (#EIACORT, Invitrogen) according to the manufacturer’s instructions. Absorbance was read at 450 nm on a fluorescence plate reader (TriStar LB941, Berthold Technologies).

### Statistical analysis

The effect size was determined based on prior experiments. Results were expressed as means ± standard error of the mean (SEM). Statistical significance was determined using parametric tests, specifically unpaired Student’s t-tests for pairwise comparisons and one- or two-way ANOVA for multiple group comparisons. Post hoc multiple comparisons were done using Sidak’s test. All datasets were screened for normality (Shapiro–Wilk test) and analysed accordingly (either ANOVA/t-tests or Mann–Whitney U test) using Prism v9.5.1 software. The criteria for significance for all analyses was set at *p < 0.05, **p < 0.01, ***p < 0.001.

## Results

### Altered GABAergic and serotonergic gene expression in the ENS of 15q duplication mice

To investigate the molecular mechanisms underlying the gastrointestinal phenotype observed in 15q dup mice, using quantitative real-time PCR (qPCR) analysis we analyzed expression of the genes in the duplicated region (Fig. 1A) in the colon, specifically focusing on the dissected smooth muscle layer and myenteric enteric neurons preparations (LMMP) (Fig. 1B). Relative to the housekeeping gene *Gapdh*, we found significantly increased expression of several GABA receptor subunit genes, including *Gabrb3*, *Gabrg3*, and *Gabra5*, in both male and female 15q dup mice compared to WT controls (Fig. 1C). Additionally, we observed elevated expression of the paternally expressed gene *Snrpn* in the 15q dup group. However, no difference was detected in the maternally expressed gene Ube3a (Fig. 1C).

**Figure 1:**
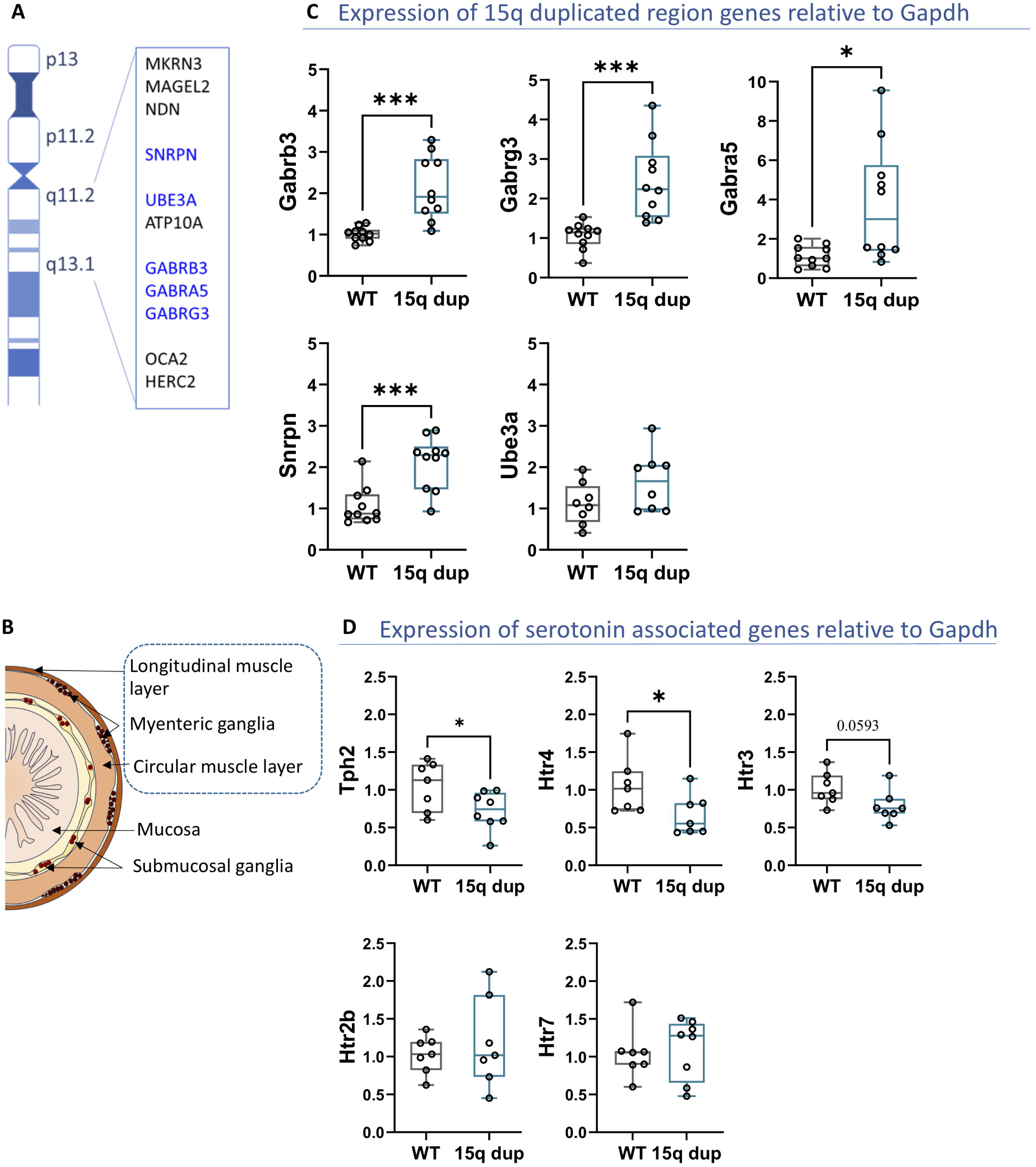
Altered GABAergic and serotonergic gene expression in the ENS of 15q dup mice. (**A**) Schematic illustrating the duplicated genes in the chromosome 15 genomic region. (**B**) Schematic of the gut cross-section indicating the LMMP preparation site from which RNA was extracted. (**C**) Expression of GABA receptor subunit genes from the Dup15q region, including *Gabrb3*, *Gabrg3*, and *Gabra5*, along with the paternally expressed gene *Snrpn* and the maternally expressed gene *Ube3a*, relative to the housekeeping gene *Gapdh*. (**D**) Relative expression of *Tph2* and 5-HT receptor genes, including *Htr4*, *Htr3*, *Htr2b*, and *Htr7*. Both male (4-5) and female (4-5) mice were used in each group. Data are represented in box plots with median and minimum and maximum of data. Student’s unpaired t-test was used to compare the means. *P<0.05, **P<0.01 and ***P<0.001.

Given that accumulating literature suggests that alterations in the GABA network can significantly impact serotonin signalling, influencing mood, behaviour, and gut-brain interactions [33], and that GABAergic activity plays a crucial role in shaping serotonin release and receptor activity by influencing the maturation and function of serotonin neurons [34–37], we investigated the impact of these potential alterations on the serotonergic system in 15q dup mice. We examined the expression of *Tph2*, the rate-limiting enzyme for serotonin biosynthesis in neurons, in RNA extracted from colon LMMP. Results revealed decreased *Tph2* expression in 15q dup mice compared to WT controls. Furthermore, we investigated the expression of 5-HT receptor genes and observed a decrease in the 5-HT4 receptor gene in 15q dup mice. Additionally, a trend towards decreased expression of the 5-HT3 receptor gene was observed in this group (p-value = 0.0593). We observed no changes to the *Tph1* gene (enzyme synthesizing the mucosal serotonin) expression in RNA extracted from the whole colon (Supplementary Fig. 1).

### 15q dup mice do not impact body weight or intestinal macroscopic structure but reveal increased crypt depths in the mid/distal colon

We observed no changes in body weight in 8–10-week-old male and female 15q dup mice. Consistent with previous findings from our group, late onset of obesity has been reported in 15q dup mice older than 20 weeks, with a significant increase in body weight observed beyond this age [38]. These weight changes were not associated with alterations in food intake, as previously reported. No overt changes were observed in cecal weight, small intestine length, or colon length in both male and female mice (Fig. 2A-B).

**Figure 2:**
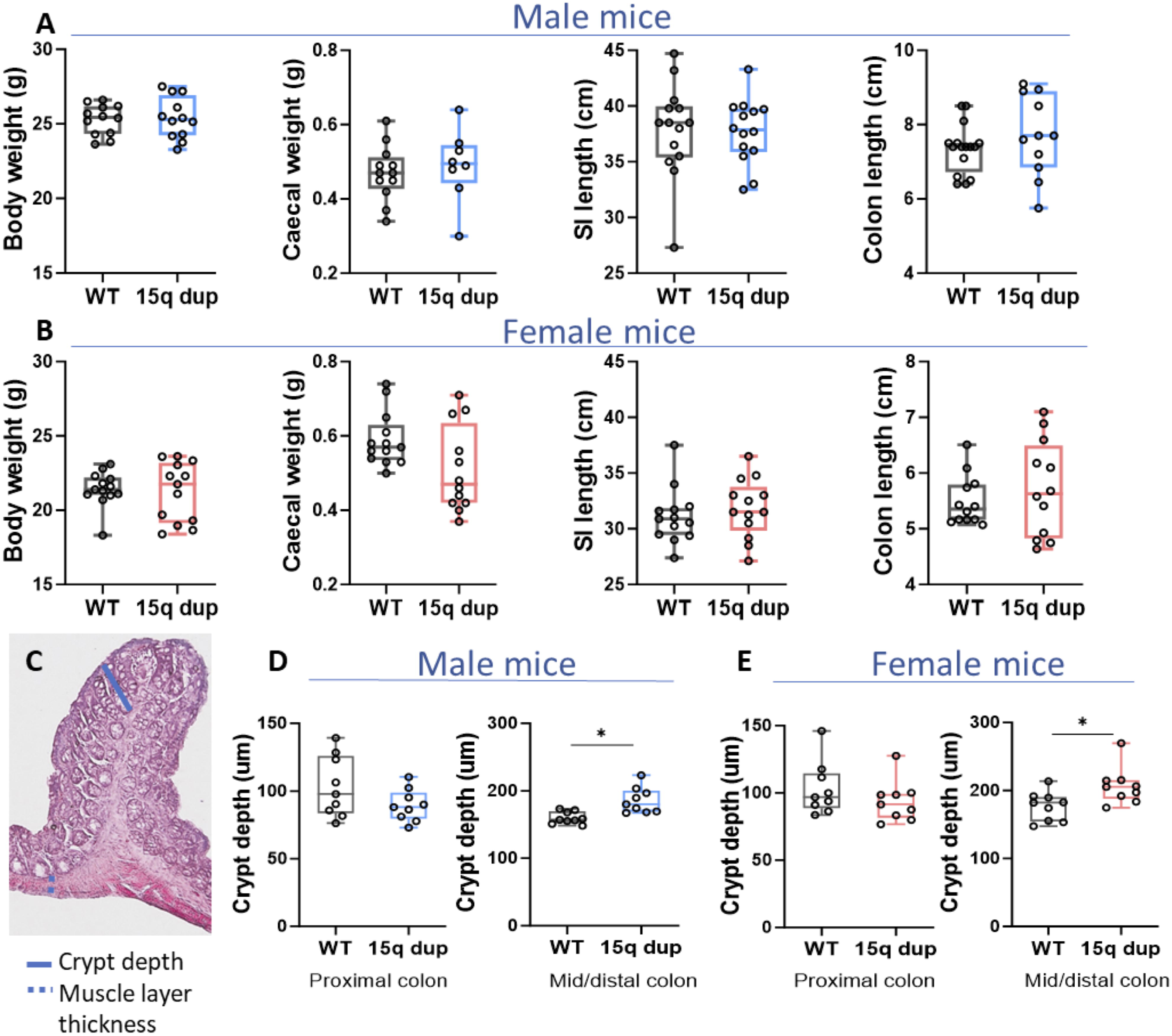
Histological analysis of the mid and distal colon in 15q dup mice reveals increased crypt depth. **(A-B)** Body weight, caecal weight, small intestinal length and colon length of male and female 15q dup mice compared to WT. **(C)** H&E-stained cross-section of the proximal colon showing the locations where crypt depth and muscle layer thickness were measured. **(D-E)** Crypt depths of proximal colon and mid/distal colon of 15q dup mice compared to WT in males and females. Data are represented in box plots with median and minimum and maximum of data. The students’ unpaired t-test was used to compare the means. Experimental groups were compared using ANOVA with repeated measures. *P<0.05

People with Dup15 syndrome often present with severe constipation-predominant gut dysfunction [8]. To investigate the underlying intestinal microscopic structure in our mouse model, we performed H&E staining (Fig. 2C). No significant changes were observed in the thickness of the muscle layer in the colon and ileum or in the villus height and width in the ileum (Supplementary Fig. 2). Both male and female 15q dup mice exhibited increased crypt depths in the mid/distal colon (Fig. 2D-E). However, in the ileum, an increase in crypt depth was observed exclusively in male mice (Supplementary Fig. 2).

### 15q dup mice lead to increased faecal pellet count, length, and reduced water content

Constipation is often characterized by infrequent bowel movements, defined as fewer than three per week [39]. Individuals with Dup15q syndrome also frequently experience constipation, which involves difficulty passing stool. Consequently, we expect an accumulation of faeces in the colon. To gain insights into the altered gut pathophysiology in 15q dup mice, we analysed their faeces. We observed a higher number of faecal pellets in the dissected colons of 15q dup mice than WT mice (Fig. 3A-C). Additionally, the length of the faecal pellets was increased in the 15q dup group for both sexes when compared to WT mice (Fig. 3D). When the percentage of water content in the faeces was measured by drying the samples overnight, the 15q dup mice exhibited a lower water content, a difference observed in both male and female mice indicating strong evidence for a constipation phenotype in 15q dup mice (Fig. 3E).

**Figure 3:**
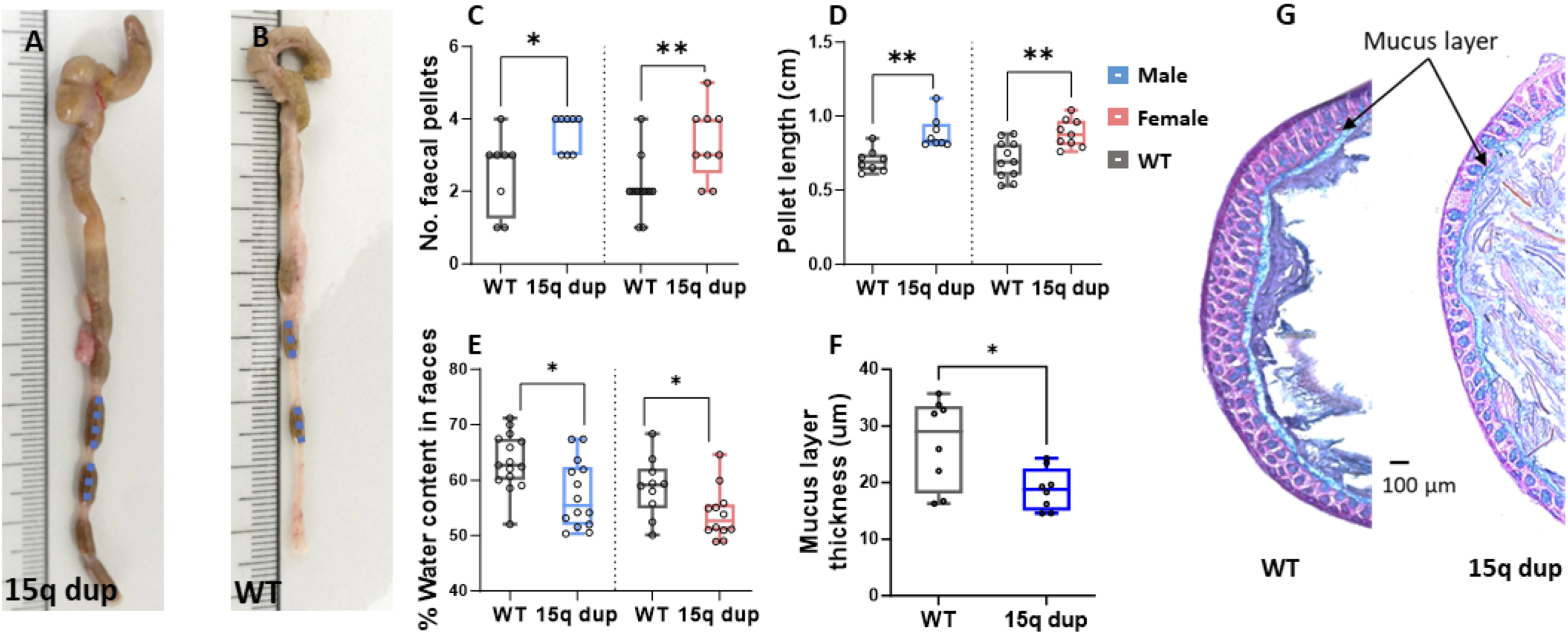
Impact of 15q dup mice on faecal characteristics. **(A-B)** Representative images of the colon from 15q dup and WT mice, respectively, showing the number of faecal pellets and their length. (**C**) Number of faecal pellets within the dissected colon of male and female 15q dup and WT mice. (**D**) Average length of faecal pellets of male and female 15q dup and WT mice. (**E**) Percentage water content of the faecal matter of WT and 15q dup mice. (**F-G**) Average mucus layer thickness in WT and 15q dup mice mid colon measured following Alcian Blue staining. Student’s unpaired t-test was used to compare the means. Experimental groups were compared using ANOVA with repeated measures. *P<0.05 and **P<0.01

Given the observed changes in faecal matter consistency and characteristics, we investigated whether the thickness of the mucus layer was altered in 15q dup mice. For this analysis, the mid-colon containing a faecal pellet was fixed, stained with Alcian Blue, and counterstained with Nuclear Fast Red. The experiment included five male and three female mice. When combining data from both sexes, we found that the thickness of the mucus layer was significantly reduced in the 15q dup group compared to WT mice (Fig. 3F-G).

### Impact of 15q duplication on the ENS and inflammatory cell populations

We performed microdissection to obtain wholemount preparations of myenteric neurons (LMMP), assessing the number of neurons per ganglion and the percentages of nitric oxide synthase (NOS), GABA, Choline Acetyltransferase (ChAT) and 5-HT-expressing neurons. There were no significant differences observed between the 15q dup and WT groups in ENS cell counts (Supplementary Fig. 3). Additionally, analysis of Iba1+ cells in colon cross-sections showed no differences between the two groups (Supplementary Fig. 3).

### Slower gut transit and altered GABA-mediated colonic motility in 15q dup mice

The pathophysiology of constipation is often multifactorial, involving disruptions in intestinal motility, secretion, absorption, permeability, sensory perception, and external factors such as diet, medications, and underlying health conditions. To investigate gut dysfunction in our model, we assessed in vivo gut motility and conducted ex vivo analyses of colonic motility and permeability. Using the intestinal sac method [40] in non-fasted mice, we found no significant differences in ileal or colonic permeability between 15q dup mice and WT controls (Supplementary Fig. 4). Since food deprivation is known to unmask changes in gut permeability, its absence in this experiment may have influenced these negative results.

Total GI transit was assessed by gavaging Carmine red dye, both male and female 15q dup mice exhibited significantly longer total GI transit times compared to WT, indicating slower GI transit in this group (Fig. 4A). Notably, we also observed slower total GI transit in female mice compared to males within WT control group, consistent with previous reports [41]. A total of 17–18 male mice and 12 female mice were tested in each group. We observed no difference in small intestinal transit between 15q dup and WT mice in both sexes (Fig. 4B). In vivo colonic transit, assessment using the glass bead propulsion assay showed 15q dup mice took significantly longer to expel the bead than WT mice, indicating slower transit (Fig. 4C).

**Figure 4:**
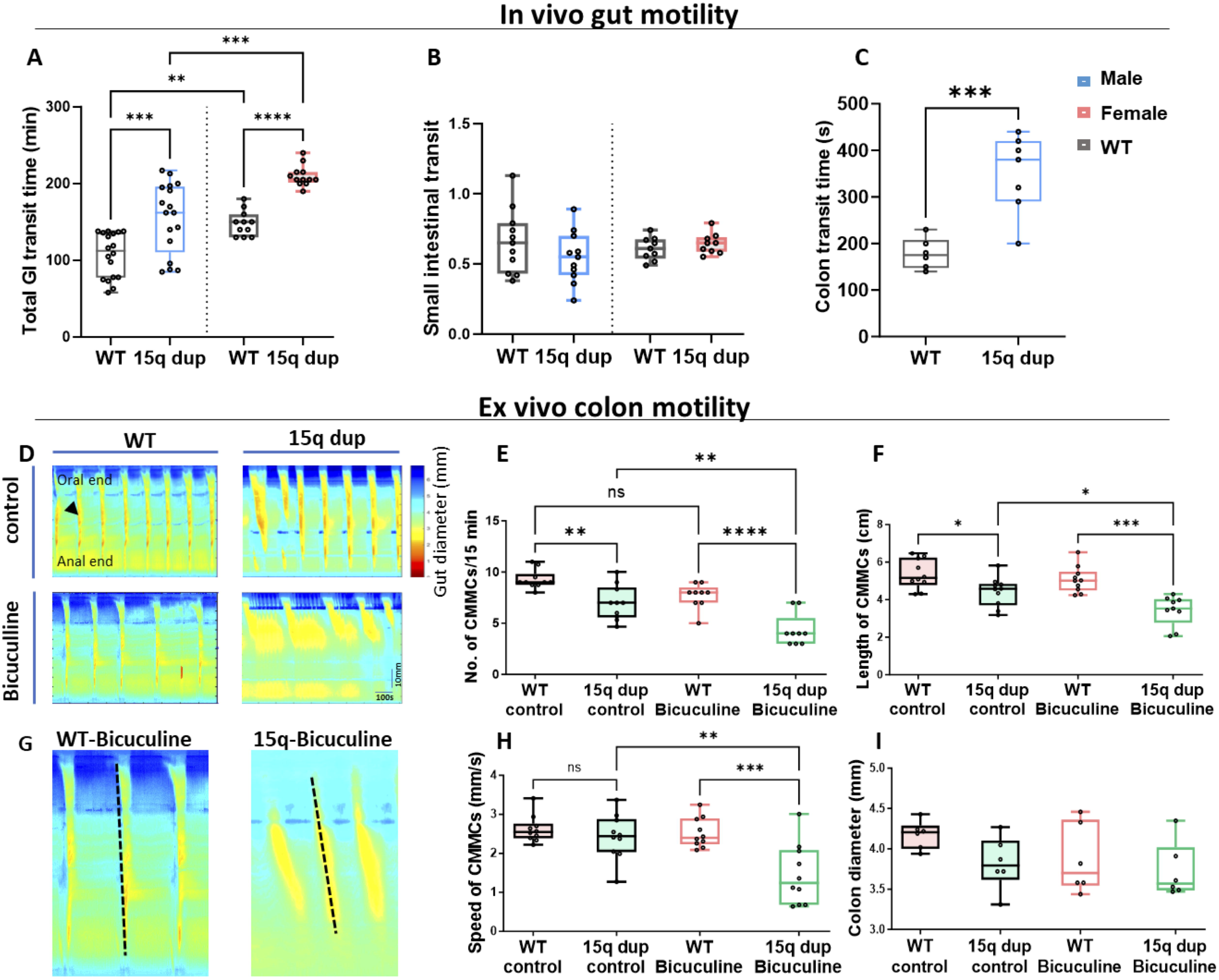
In vivo and Ex vivo analysis of intestinal motility of 15q dup mice. (**A**) In vivo total gastrointestinal (GI) transit time in male and female 15q dup mice was compared with respective WT mice. (**B**) Small intestinal motility is represented by a ratio of dye front to the small intestine/length of the small intestine. (**C**) In vivo colonic transit time was measured in male mice compared with WT mice. (**D-I**) Ex vivo colonic motility was assed analysing colonic migrating motor complexes (CMMCs) in male mice. (**D**) Representative spatiotemporal maps where the diameter of the colon is plotted against the length of the tissue as a function of time (arrowhead indicates a representative CMMC). Detailed CMMC characteristics such as (**E**) count for 15 min, (**F**) length in cm, (**G-H**) speed of contractions (mm/s) analysed using the slope of CMMCs (**I**) and resting colon diameter (mm) were compared against WT. The students’ unpaired t-test was used to compare the means. Experimental groups were compared using ANOVA with repeated measures. *P<0.05, **P<0.01 and ***P<0.001

To further investigate colonic motility, we employed ex vivo video imaging of gut motility (Fig. 4D) and we measured and compared the frequency, length, and speed of contractions, as well as the resting colon diameter, between WT and 15q dup mice under control conditions and following exposure to the GABAA receptor antagonist Bicuculline. Under control conditions, where colons were exposed to saline solution, 15q dup male mice exhibited fewer colonic migrating motor complexes (CMMCs) than WT mice. Bicuculline exposure further reduced the CMMC frequency in the 15q dup group relative to WT (Fig. 4D-E). Bicuculline did not alter CMMC frequency in the WT colon (P = 0.071), consistent with previous findings [14]. During saline exposure, 15q dup mice exhibited shorter contractions compared to WT mice, and the contraction length was further reduced during Bicuculline exposure. In contrast, Bicuculline did not alter CMMC length in WT mice (Fig. 4F). The speed of contractions (Fig. 4G-H) and resting gut diameter (Fig. 4I) were not different between 15q dup and WT mice under control conditions. Upon exposure to Bicuculline, the speed of contractions was reduced in the 15q dup group, resulting in much slower CMMCs, while no effect was observed in the WT group (Fig. 4H). Resting colon diameter was unaffected by Bicuculline in both WT and 15q dup mice (Fig. 4I).

### Prucalopride (5-HT4R agonist) restores altered gastrointestinal motility and improves social behaviour in 15q dup mice

This study identified GABAA receptor targeting as a potential approach for treating constipation in our model, though systemic exposure poses risks like seizures. Given serotonin’s role in gut motility, we explored serotonin receptor modulation and found reduced Tph2 and Htr4 expression, indicating altered serotonin signalling. To further investigate this, we focused on targeting the 5HT4R, hypothesizing that modulation of this receptor could alleviate the gut dysfunction observed. Prucalopride, a 5-HT4R agonist approved by the FDA for treating chronic constipation, was chosen for this purpose. Following Prucalopride administration via oral gavage for 6 days, we assessed several key parameters: total GI transit, body weight, reciprocal social interaction, and corticosterone concentration in faeces (Fig. 5A). Sham control mice were treated with an equal volume of saline for comparison. We presented the data in the male mice to eliminate the potential confounding effects of cyclic changes in gut motility associated with the estrus cycle in female mice, as we previously reported [29, 42].

**Figure 5:**
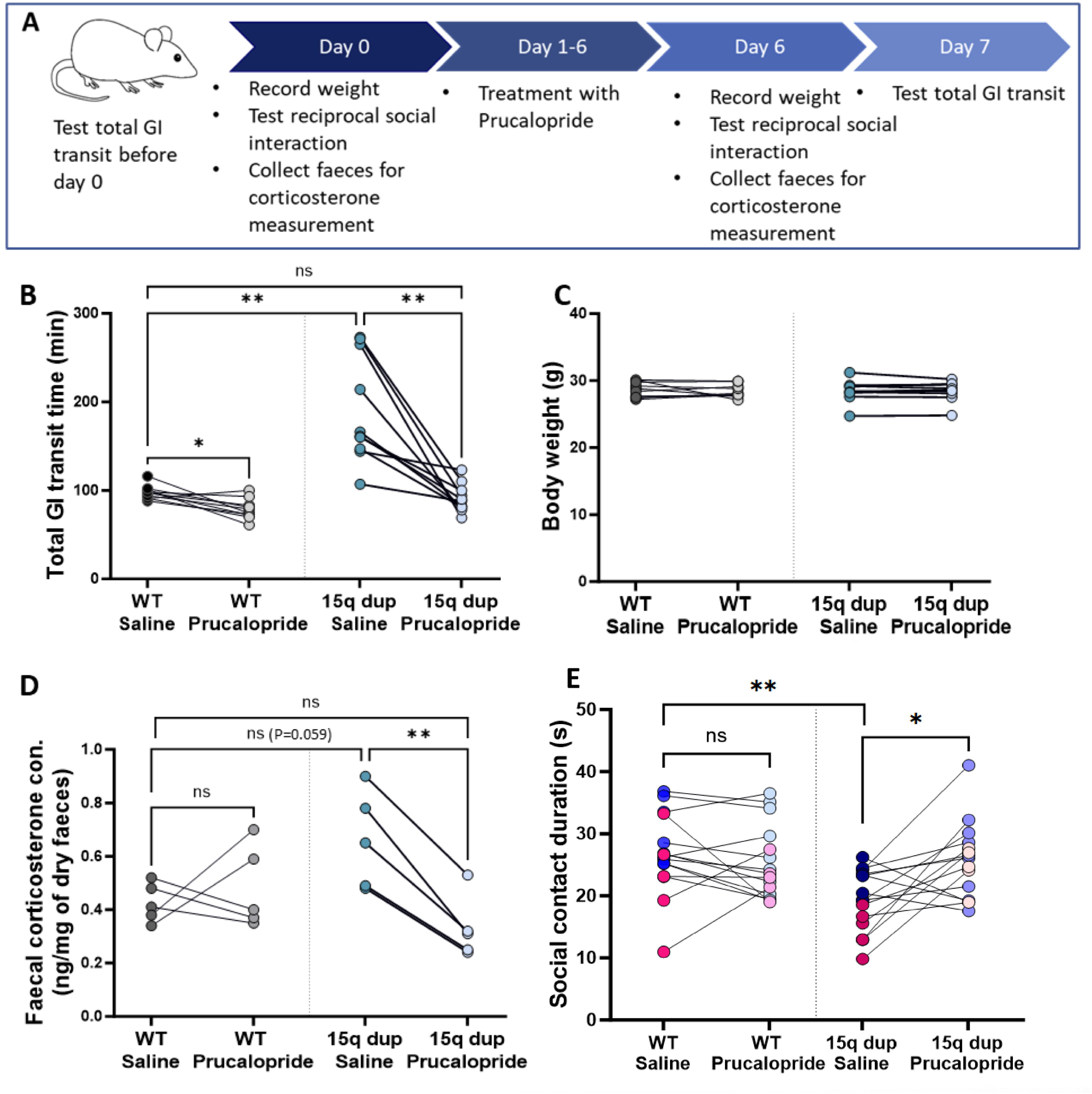
Effects of Prucalopride (5-HT4R agonist) on GI motility and social behaviour on 15q dup mice. **(A)** Schematic representation of the Prucalopride treatment experimental plan. **(B)** Pairwise comparison of the total GI transit times between saline and Prucalopride treatment. **(C)** Pairwise comparison of the body weight change during the 6-day treatment period**. (D)** Pairwise comparison of the faecal corticosterone concentration between saline and Prucalopride treatment. **(E)** Social contact duration between saline and Prucalopride treatment during reciprocal social interaction test. Data from males are presented in blue and female in pink. Student’s unpaired t-test was used to compare the means. Experimental groups were compared using ANOVA with repeated measures. *P<0.05 and **P<0.01

We first assessed the baseline GI transit in saline treated sham controls which was slower in 15q dup mice compared to WT (Fig. 4A and 5B). Treatment with Prucalopride significantly reduced GI transit time in the 15q dup group, restoring it to levels comparable to WT mice (Fig. 5B). This suggests that Prucalopride effectively alleviates the delayed GI transit observed in 15q dup mice. Notably, Prucalopride also reduced GI transit time in WT mice, which is consistent with the known effects of the drug on GI motility [43]. No change in body weight was observed in either group during treatment (Fig. 5C).

Patient surveys in Dup15q syndrome have reported a correlation between GI dysfunction and challenging behaviours, including aggression [8, 9]. Moreover, successful treatment of GI issues has been linked to improvements in these behaviours [8]. Building on these findings, we sought to investigate whether our treatment could positively impact reciprocal social interaction and stress-related corticosterone levels in faeces collected during the reciprocal social interaction test. Faecal corticosterone was assessed as a non-invasive biomarker of chronic stress [44], as elevated levels have been linked to gut dysfunction, behavioural abnormalities, and aggression in neurodevelopmental disorders [45]. Each experimental group consisted of 24 mice. Treatment with Prucalopride significantly reduced faecal corticosterone levels in 15q dup mice compared to saline-treated sham control mice, bringing the levels down to those observed in WT mice (Fig. 5D). Each data point in the graph represents the average faecal analysis from two individual mice. In the reciprocal social interaction test, at baseline, saline-treated 15q dup mice showed reduced social contact duration compared to WT mice, as previously reported [20, 46] (Fig. 5E). We report that the treatment with Prucalopride significantly increased social contact duration in 15q dup mice relative to the saline-treated 15q dup group, with no effect on WT mice (Fig. 5E) and no sex-specific differences. The treatment notably improved social contact duration in 15q dup mice, bringing it in line with WT levels (Fig. 5E).

## Discussion

In this study, we investigate GI dysfunction in a mouse model of Dup15q (15q dup), a validated model for neurodevelopmental disorders, including autism [15, 20–22, 46]. Our findings reveal a significant alteration in GI motility in 15q dup mice, which may contribute to the constipation-predominant gut dysfunction commonly observed in individuals with Dup15q syndrome. Through molecular analysis, we identify disruptions in both GABAergic and serotonergic signalling pathways as key factors underlying these GI disturbances. Importantly, we demonstrate that Prucalopride, a 5-HT4R agonist, effectively restores GI dysmotility and improves social behaviour in 15q dup mice, highlighting its potential as a therapeutic strategy for managing both gastrointestinal and behavioural challenges in this model.

### GI dysfunction in 15q dup mice: A novel discovery

In this study, we report, for the first time, significant GI dysfunction in the Dup15q mouse model. This discovery highlights the relevance of this model for studying the link between neurodevelopmental disorders, such as ASD, and gut dysfunction, which is commonly observed in affected individuals. Our findings provide valuable insights into the gut motility disturbances, altered faecal characteristics, and molecular mechanisms underlying the GI challenges experienced by individuals with Dup15q syndrome.

We identified disruptions in the GABAergic and serotonergic systems as key contributors to GI dysfunction in 15q dup mice. Specifically, we observed increased expression of GABA receptor subunits in both male and female mice, indicating an altered GABAergic signalling network. This is consistent with recent studies suggesting that GABAergic dysregulation may influence various physiological processes, including serotonin signalling [33, 36]. RNA-sequencing data on ENS gene expression, in conjunction with immunofluorescence and transgenic mouse studies by Morarach et al. (2020), reveals that ENS cell subclass 12, which includes ligands for Tph2, also expresses several key GABA receptor subunits, including Gabrb3, Gabrg2, Gabrg3, and Gabbr1 [47]. This highlights the potential for GABAergic modulation in Tph2-positive enteric neurons, suggesting a functional interplay between GABA and serotonin signalling in this subset of ENS cells. Additionally, we found reduced expression of serotonin-related genes, such as Tph2 and Htr4, in a preparation containing enteric neurons. This suggests that altered GABAergic signalling may disrupt serotonergic neurotransmission, which is crucial for regulating mood, behaviour, and gut motility [36]. Therefore, the combined dysregulation of both GABA and serotonin likely contributes to the observed GI dysfunction in 15q dup mice. The findings also highlight the importance of neuronal serotonin in mediating normal gut motility.

Our motility assays further demonstrated that 15q dup mice exhibit slower gastrointestinal transit, consistent with the constipation observed in this model. Ex vivo studies confirmed this finding, showing a reduction in colonic motility and a significant decrease in the frequency of CMMCs. These motility changes were mediated by the GABAergic pathway, as indicated by the further reduction in motility following exposure to the GABAA receptor antagonist Bicuculline. Importantly, no significant differences in small intestinal motility were observed, suggesting that the motility deficits are more pronounced in the colon, where GABA and serotonin play a particularly strong role in regulating peristalsis and contractile activity [48–52]. Alterations to the GABA and serotonin-associated colonic motility changes were observed in other preclinical autism mouse models [13, 14, 16, 53]. These findings underscore the importance of GABAergic and serotonergic systems in maintaining gut motility and suggest that their dysregulation may be central to the GI dysfunction observed in Dup15q syndrome.

### Prucalopride: A dual therapeutic approach to restore gut dysmotility and improve behaviour

Prucalopride, a highly selective 5-HT4R agonist, emerged as a promising therapeutic intervention in our model, restoring normal gastrointestinal motility and improving social behaviour. The use of Prucalopride treatment in 15q dup mice significantly reduced GI transit time, bringing it to levels comparable to WT mice, and alleviated the constipation-associated gut dysfunction. Importantly, Prucalopride also improved reciprocal social interaction and reduced corticosterone levels in faeces, suggesting a beneficial impact on both GI function and behaviour. This result aligns with emerging literature highlighting the therapeutic potential of 5-HT4R agonists for managing gut dysfunction in neurodevelopmental disorders. Prucalopride has been recommended by both the US Food and Drug Administration and the European Union for the symptomatic treatment of chronic idiopathic constipation (CIC), particularly in patients who have not experienced sufficient symptom relief from common laxatives [18]. Studies have shown that Prucalopride 1-2 mg dose can effectively treat constipation-predominant disorders by enhancing gut motility and improving overall gastrointestinal health, including CIC [19, 43]. Furthermore, Prucalopride has been shown to improve memory and cognition in both animal models and humans [54–56]. Consistent with previous studies suggesting that alleviating GI dysfunction can reduce behavioural challenges, such as aggression and irritability, in Dup15q syndrome [8, 9], our findings that Prucalopride improves social behaviour align with and are supported by this body of literature. This also implicates a potential relationship between gut-brain interactions and social behaviour in this model. Our findings suggest that the Prucalopride treatment may be linked to reductions in stress-related corticosterone levels. The HPA (hypothalamic-pituitary-adrenal) axis is known to be involved in the gut-brain connection, with stress and anxiety having a direct impact on gut motility and permeability [57]. 5HT4R are directly associated with stress levels [58] and 5HT4R agonists are reported to hold anxiolytic effects [59]. In our study, Prucalopride treatment reduced faecal corticosterone levels in 15q dup mice, which may reflect reduced stress and anxiety levels following improved gastrointestinal function or systemic effects on 5HT4R in the brain. Prucalopride has emerged as a promising therapeutic agent with multiple beneficial effects on the enteric nervous system (ENS). It has been shown to support ENS development by promoting neurogenesis and neuroprotection [60–62], enhance mucus secretion from goblet cells [50, 63] and stimulate acetylcholine release, thereby improving neurotransmission [64]. Additionally, Prucalopride exhibits anti-inflammatory properties [32], further highlighting its potential as a multifaceted treatment for gastrointestinal and neurodevelopmental disorders.

The study highlights potential treatments for GI dysfunction in individuals with Dup15q syndrome, who frequently suffer from symptoms like constipation, irritability, and aggression. Alterations in GABAergic and serotonergic signalling were identified as contributors to these dysfunctions in the Dup15q mouse model. Prucalopride, a 5-HT4R agonist, improved both GI motility and social behaviour in these mice, suggesting its therapeutic potential for addressing both physical and behavioural symptoms. These findings support the need for clinical trials to evaluate Prucalopride’s dual benefits for Dup15q syndrome.

While this study revealed important insights, several limitations remain. The gut-brain axis’s role in cognition and stress was not fully explored, and other behavioural assessments, such as cognitive tests, were not included. The role of 5-HT3 receptors in GI dysfunction remains unclear, as their reduced expression did not reach statistical significance. A recent report proposed a drug to inhibit serotonin transporter in the intestinal epithelium could be an effective agent for both GI and mood dysfunctions [2]. Additionally, investigating GABA-A receptor modulators like Basmisanil [65], could offer promising therapeutic options. The study used a paternal Dup15q model, which, though validated for studying associated phenotypes [21], does not fully replicate the maternal duplication most commonly linked to the syndrome [66]. Future studies should validate findings in maternal duplication models.

In conclusion, we present the Dup15q mouse model as a valuable tool for studying gut dysfunction in neurodevelopmental disorders such as autism. The underlying mechanisms of gastrointestinal dysfunction in this model involve alterations in GABAergic and serotonergic signalling, leading to disrupted gut motility indicating constipation. Our study highlights the potential of Prucalopride, a 5-HT4R agonist, to restore gut dysmotility and improve behavioural challenges. Given the emerging evidence supporting the role of serotonin in regulating both gut function and behaviour, Prucalopride may be a promising therapeutic strategy for improving the quality of life in individuals with Dup15q syndrome and similar neurodevelopmental disorders.

## Disclosure statement

All authors declare no competing financial or personal interests that could have influenced the findings in this paper.

## Grants

Japanese Society for Promotion of Science (JSPS) Postdoctoral Fellowship (P20704) to GKB; JSPS KAKENHI (16H06316, 20F20704, 21H04813, 23KK0132, 23H04233, 24K22036, 24H00620, 24H01241) to TT, (23K06399) to KT; Japan Agency for Medical Research and Development (AMED), JP21wm0425011 to TT; Japan Science and Technology Agency (JST), JPMJPF2018, JPMJMS2299, JPMJMS229B to TT; Intramural Research Grant (6-9) for Neurological and Psychiatric Disorders of NCNP to TT; Takeda Science Foundation to KT and TT; Taiju Life Social Welfare Foundation to TT; Kobe University Strategic International Collaboration Research Grant 2023/24 for GKB and TT

## CRediT Authorship Contributions

Balasuriya, G.K., PhD (Conceptualization, Data curation, Formal analysis, Funding acquisition, Investigation, Methodology, Validation, Visualization, Writing – review & editing)

Tamada, K., PhD (Data curation, Methodology, Validation, Visualization, Review & editing)

Nomura, J., PhD (Data curation, Methodology, Validation, Visualization)

Cirillo, C., PhD (Data curation, Methodology, Review & editing)

Takumi, T., MD, PhD (Conceptualization, Resources, Supervision, Funding acquisition, Methodology, Review & editing)

## Data and Materials Availability

All data supporting the findings of this study are provided within the main text or the supplementary information. Raw datasets and in-house software used for data collection and analysis are available upon reasonable request.

## Acknowledgments

We thank Yuko Tamai, Hiroko Maegawa, Hiroko Okubo, and all technical staff in the Takumi lab for technical assistance.

## Notes

### Competing Interest Statement

The authors have declared no competing interest.

