## supplemental info for "Prucalopride ameliorates delayed gastrointestinal transit and social behaviour in a mouse model of 15q duplication syndrome": Supplementary info.docx

**Supplementary materials**

**Supplementary Table 1**: Forward (Fw.) and Reverse (Rv.) primer sequences used for quantitative PCR reactions

| Gene | Strain | Sequence#1 Fw. | Sequence#2 Rv. |
| --- | --- | --- | --- |
| Gapdh | m | ACGGGAAGCTCACTGGCATGGCCTT | CATGAGGTCCACCACCCTGTTGCTG |
| Snrpn | m | GCAAAACAGCCAGAACGTGAA | GCACACGAGCAATGCCAGTAT |
| Ube3a | m | TCTGCTGCTGCTATGGAAGA | CACATTCCACGTTAGGTGACA |
| Gabrb3 | m | GAATGTTGTCTTCGCCACAGGT | ACCCACGAGAGGATTGTGATCA |
| Gabra5 | m | TCCAAACATCCCAAAAGAGC | GAGAGGTGGCCCCTTTTATC |
| Gabrg3 | m | CGAATAAGCCTTCAAGCACCC | AGGTGTCCTCAAATTCCTGCC |
| Tph2 | m | TTCGTCCATCGGAGAATTGAAG | GCGTCCTGAAAGGTGGTGATTA |
| Tph1 | m | AACAAAGACCATTCCTCCGAAAG | TGTAACAGGCTCACATGATTCTC |
| Htr4 | m | GCTAATGTGAGT TCCAACGA | GGTAAGTAGGACATCCAGAG |
| Htr3 | m | ATCAATGAGTTTGTGGACGTG | GAAGATGCTCTTGTCAGACC |
| Ht2rb | m | ATCATGTTTGAGGCTATATGGC | CACTGATTGGCCTGAATTGG |
| Hhtr7 | m | GTTAGTGTCACGGACCTCAT | ATCATTTTGGCCATACATTT |

**Supplementary Table 2**: Details of the primary and secondary antibody used for the immunofluorescence experiments

| Antibody | Host species | Source | dilution |
| --- | --- | --- | --- |
| Hu C/D | Mouse | Invitrogen #A21271 | 1 in 500 |
| nNOS | Goat | abcam #AB1376 | 1 in 500 |
| GABA | Rabbit | Sigma #A2052 | 1 in 500 |
| 5-HT | Rabbit | ImmunoStar #20080 | 1 in 500 |
| ChAT | Goat | Chemicon #AB144P | 1 in 250 |
| Iba1 | Rabbit | Wako #019-19741 | 1 in 500 |
| Alexa Fluoro anti goat 647 | Donkey | abcam #ab150131 | 1 in 400 |
| Alexa Fluoro anti mouse 568 | Donkey | Invitrogen #A10037 | 1 in 500 |
| Alexa Fluoro anti rabbit 488 | Donkey | Invitrogen #A21206 | 1 in 500 |

**Supplementary methods**:

**Video Imaging technique to assess Ex vivo colonic motility**

Adult male WT and 15q dup mice (aged 8–10 weeks) were culled by cervical dislocation and the entire colon was dissected out to study the ex vivo colonic motility using video imaging technique as previously reported [28, 29]. Briefly, the dissected colon was mounted on an organ bath superfused with Kreb’s saline solution bubbled with carbogen gas course (Kreb’s composition, mM: 118 NaCl, 4.76 KCl, 0.99 NaH2PO4.2H2O, 1.20 MgSO4.7H2O, 2.5 CaCl2.2H2O, 11.1 D-glucose, 25 NaHCO3) and the temperature of the organ bath was maintained between 34 - 37 °C. The oral end of the colon was connected to a Kreb’s solution supply that maintained an intraluminal pressure of 2-4 cm H2O. Spontaneous contractions of the colon were video recorded using Logitech Quickcam Pro webcam mounting it right above the organ bath. Recorded videos were then processed through an in-house edge detection software (Scribble 2.0) to generate spatiotemporal maps reflecting the colonic motility patterns. Spatiotemporal maps were analysed using purpose-built MATLAB plugin, Analyse 2.0.

**Alcian Blue staining for mucus layer thickness**

Alcian Blue staining was performed on mid-colon tissues (1–2 cm segments containing a faecal pellet) following an established protocol Tissues were fixed in Carnoy's fixative overnight, washed with PBS, and cryoprotected in a 30% sucrose solution. Subsequently, samples were embedded in Optimal Cutting Temperature (OCT, Tissue Tek.) compound using cryomolds and sectioned at a thickness of 10 µm with a cryostat. Tissue sections mounted on slides were washed with distilled water and incubated in 30% acetic acid for 30 minutes to enhance staining. Sections were then stained with Alcian Blue solution (Sigma) for 30 minutes, followed by rinsing under running tap water for 1 minute. Counterstaining was performed with Nuclear Fast Red solution (Sigma) for 30 minutes. After thorough washing, the slides were mounted using a standard mounting medium. Brightfield images of the stained sections were acquired using an Olympus slide scanner (Olympus VS120). The mucus layer thickness was quantified using ImageJ software (NIH, USA), and the average thickness was calculated for each sample.

**Supplementary Figure 1**

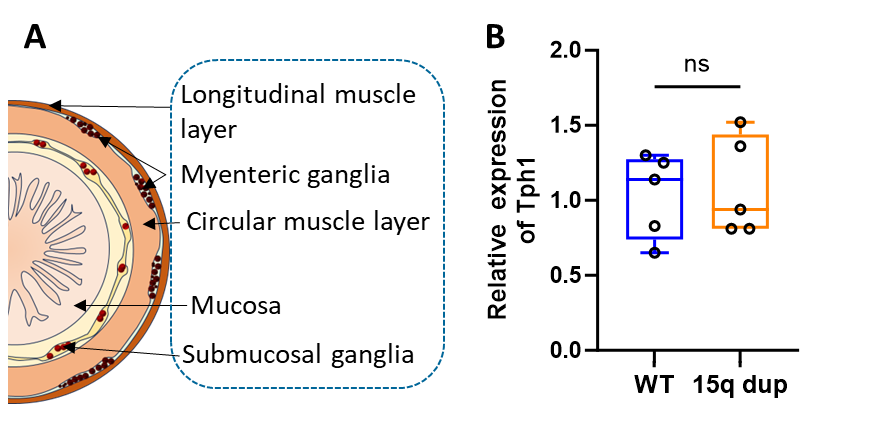

**Supplementary Figure 1: Tph1 gene expression in the colon and histological analysis of the small intestine. (A)** Schematic cross section of the colon indicating all the layers from which the RNA was extracted and (**B)** the relative expression of Tph1 was compared between Dup15q and WT. Data are represented in box plots with median and minimum and maximum of data. The students’ unpaired t-test was used to compare the means.

**Supplementary Figure 2**

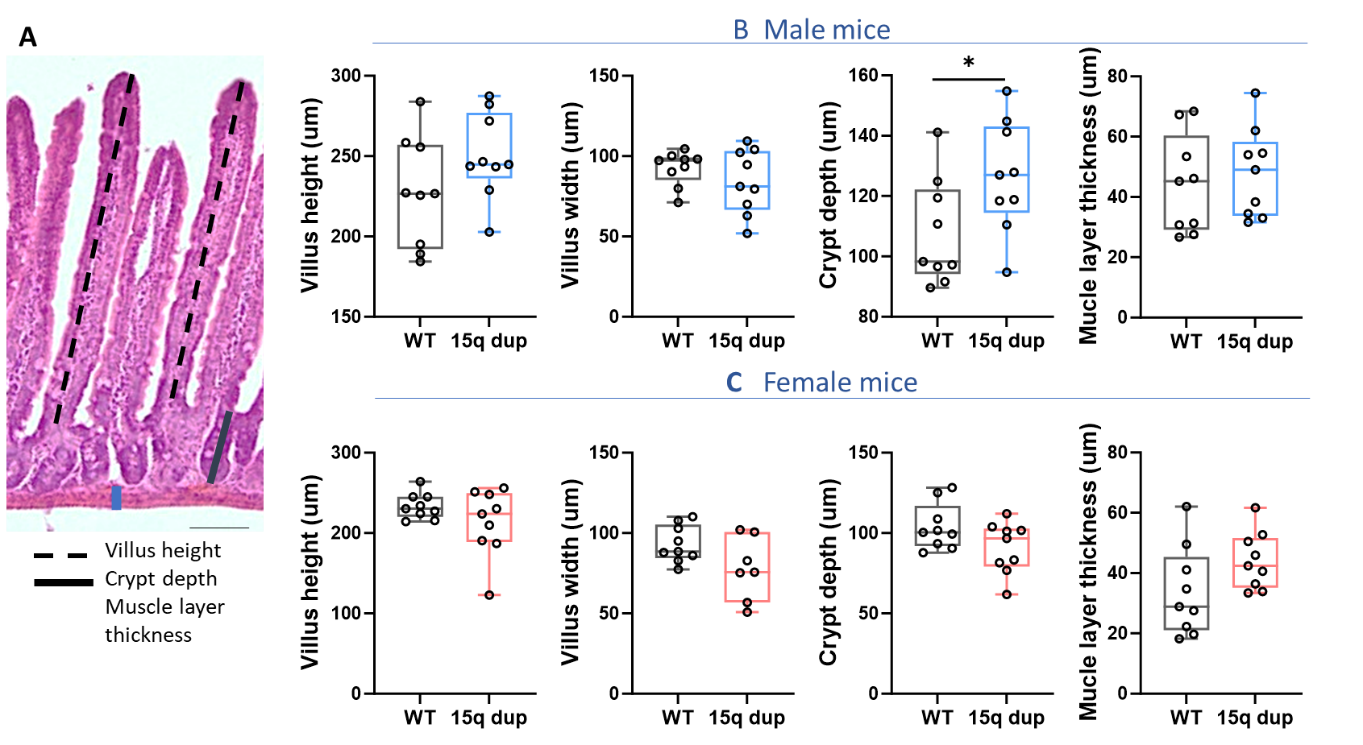

**Supplementary Figure 2: Histological analysis of the small intestine. (A)** H&E-stained cross section of the ileum illustrating where measurements were made for the villus height, crypt depth and the muscle layer thickness. **(B-C)** Average villus height, villus width, crypt depth and the muscle layer thickness of male and female 15q dup mice compared to WT. Data are represented in box plots with median and minimum and maximum of data. The students’ unpaired t-test was used to compare the means. Experimental groups were compared using ANOVA with repeated measures. *P<0.05

**Supplementary Figure 3**

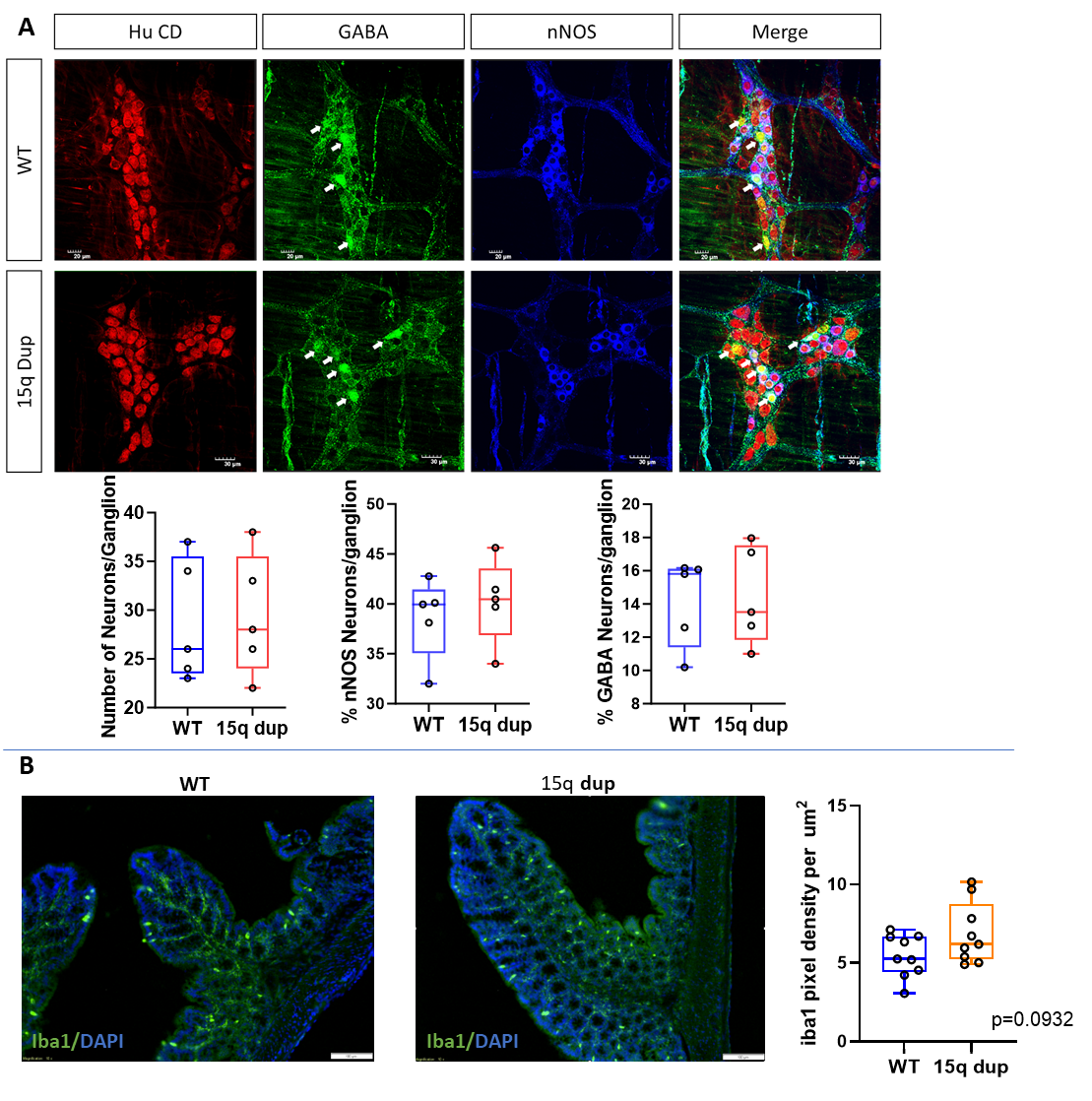

**Supplementary Figure 3: Enteric nervous system arrangement in the colon and Iba1^+^ cells in the colon mucosa. (A)** Representative images showing immunofluorescence labelling of pan-neuronal marker HuC/D, GABA, and nNOS in whole-mount preparations of the proximal colon from 15q dup and WT mice. **(B)** Mid colon cross sections immune labelled with Iba1 and DAPI. Data are represented in box plots with median and minimum and maximum of data. The students’ unpaired t-test was used to compare the means.

**Supplementary Figure 4**

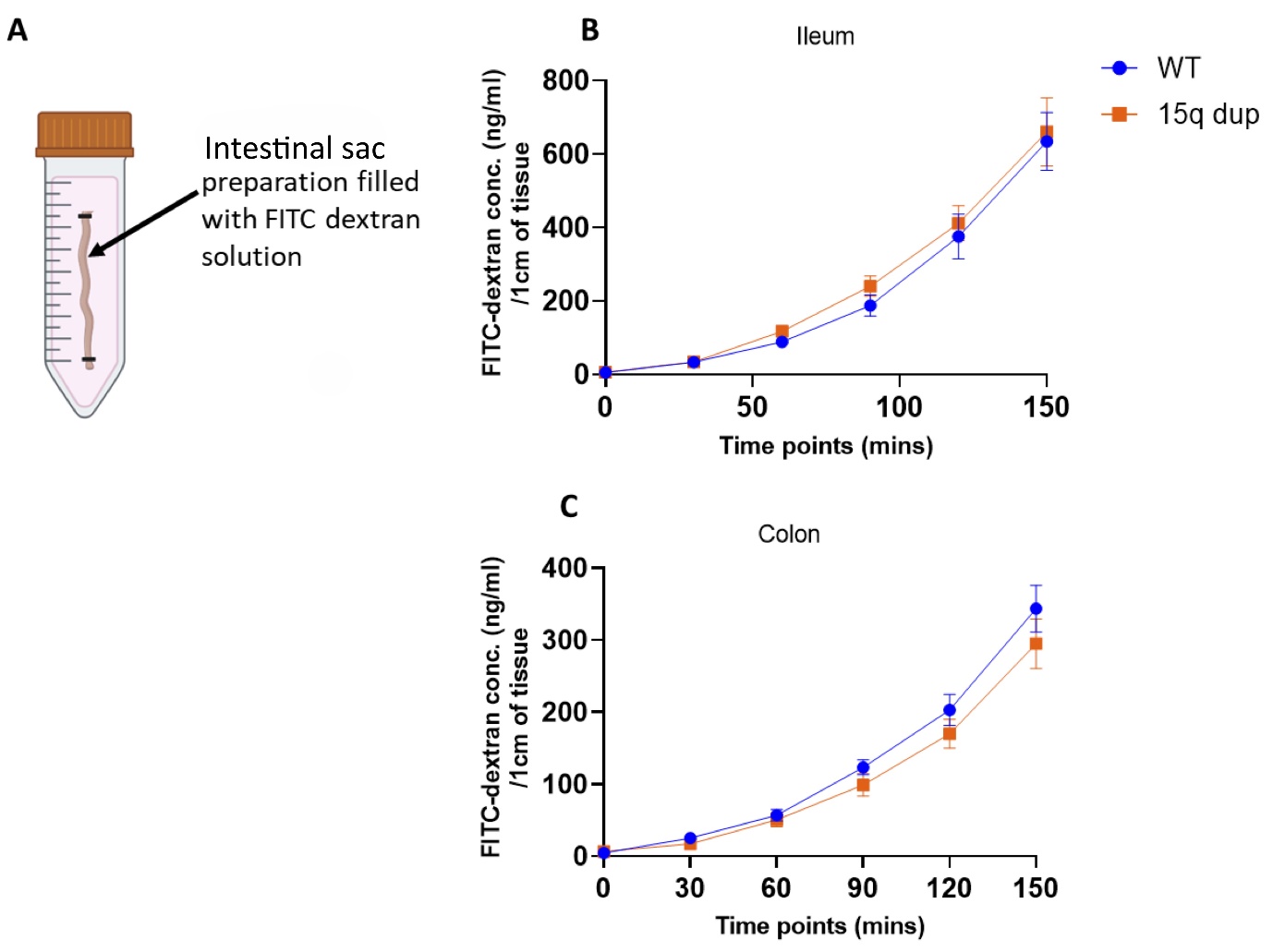

**Supplementary Figure 4: Intestinal permeability measured using the intestinal sac method. (A)** Diagram representing the gut sausage preparation filled with FITC tagged dextran solution in DMEM F/12 solution during permeability measurement. (**B-C**) FITC-dextran concentrations in the ileum and colon of WT and 15q dup mice were measured over a 150-minute time period. The students' unpaired t-test was used to compare the means.
