## Supplementary figures and images for "Prucalopride ameliorates delayed gastrointestinal transit and social behaviour in a mouse model of 15q duplication syndrome"

### figures_S1.tiff

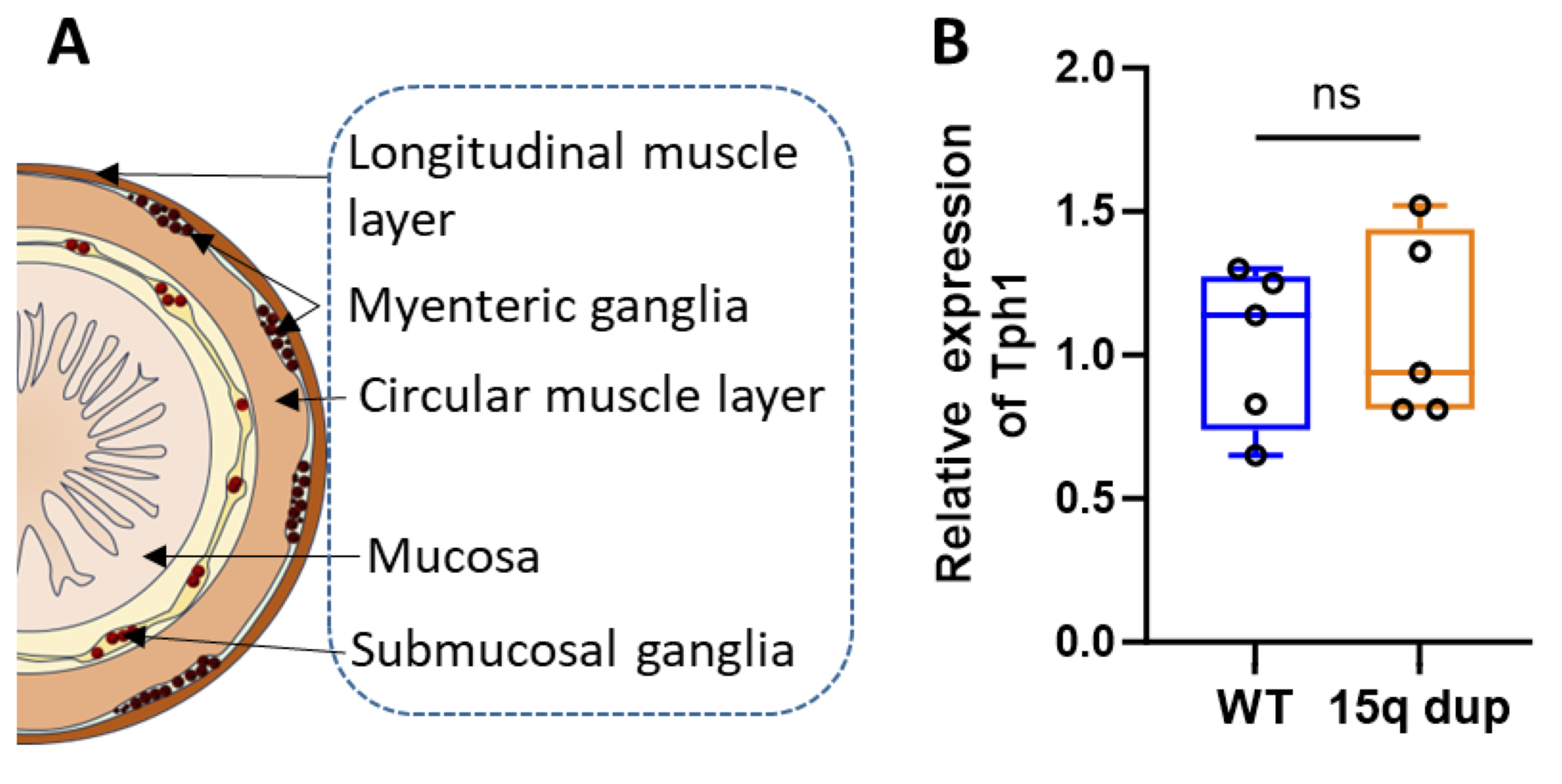

### figures_S2.tiff

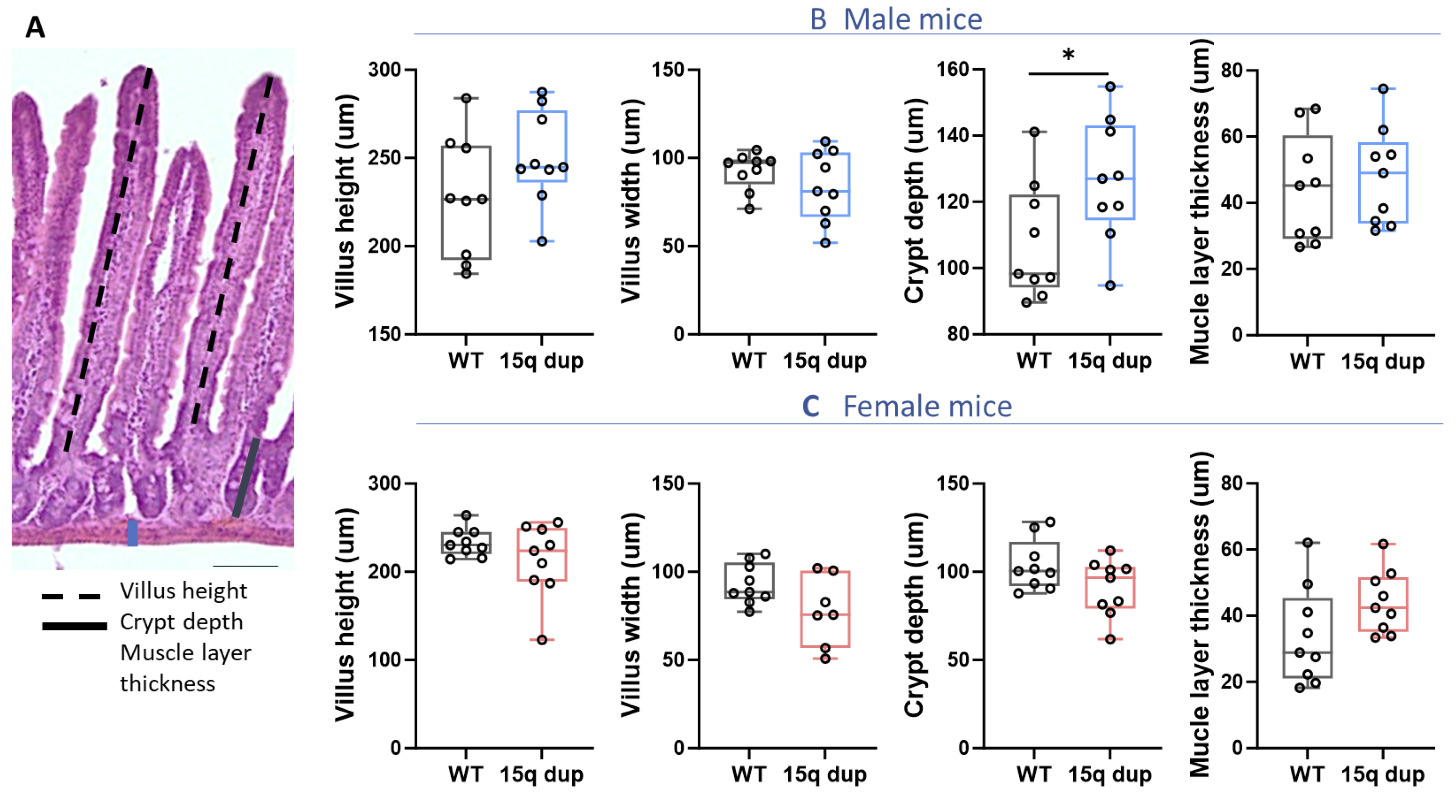

### figures_S3.tiff

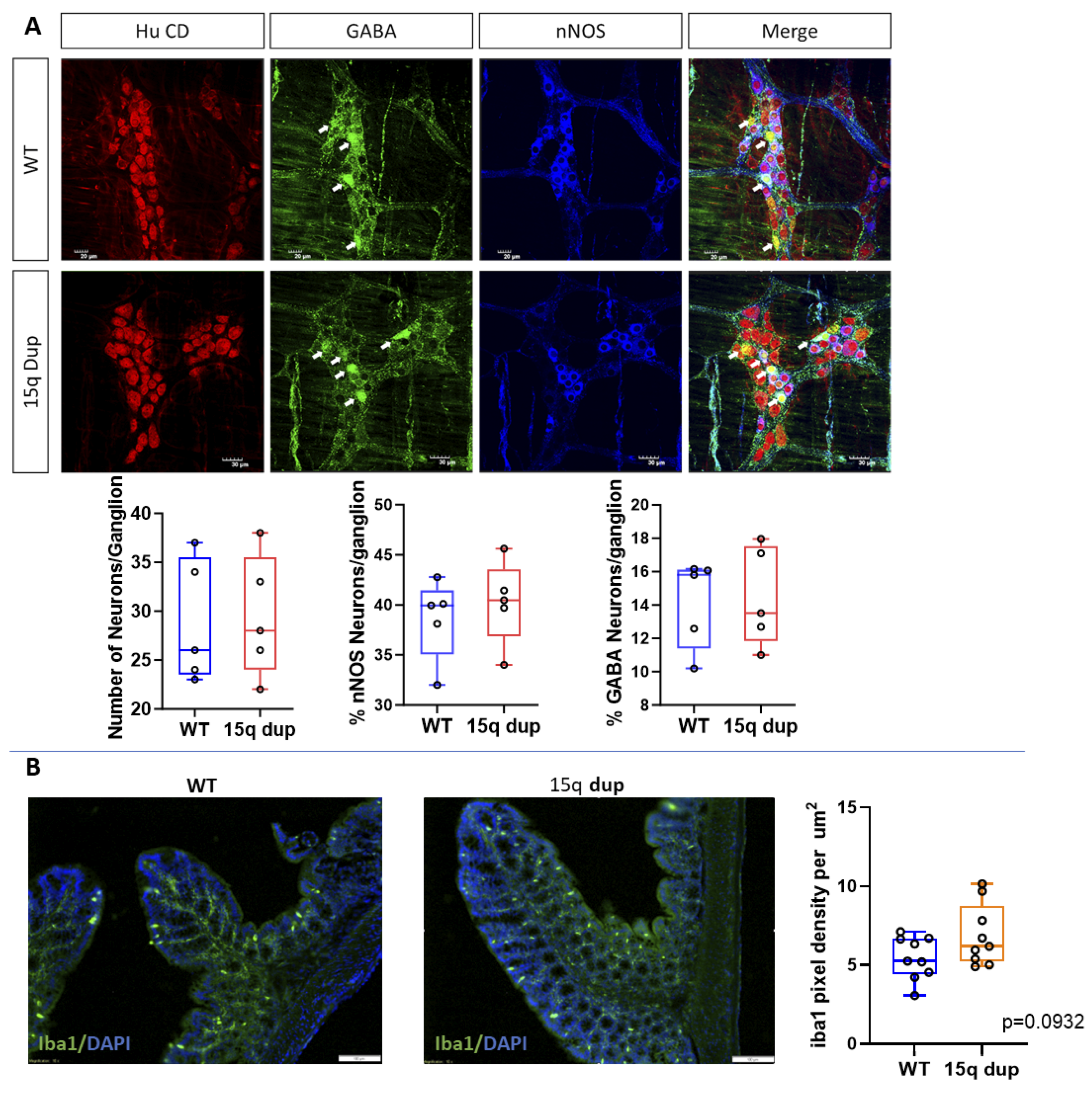

### figures_S4-new.tiff

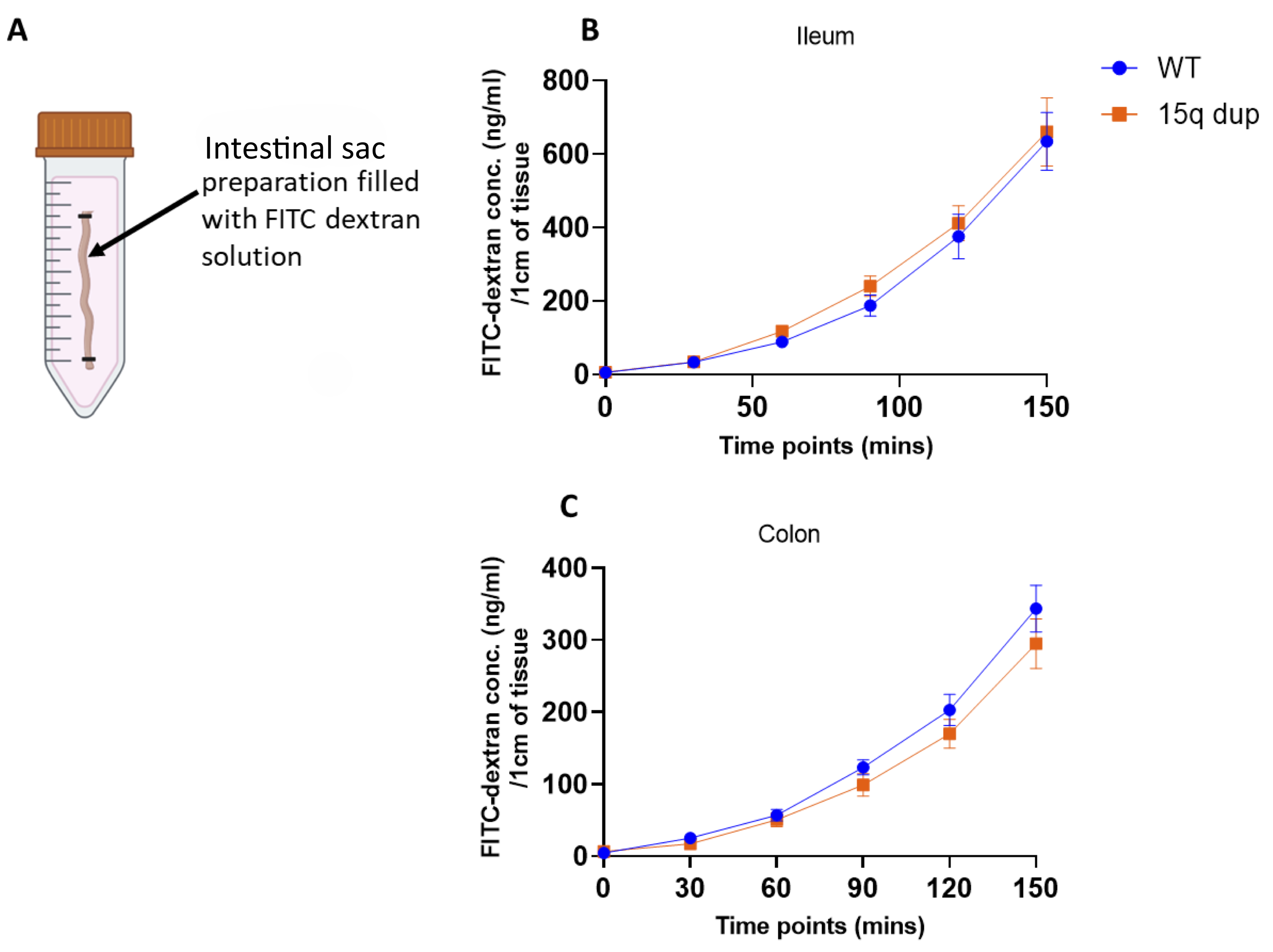
